# Polyunsaturated fatty acid-derived lipid mediator networks characterize COVID-19 severity and risk for critical illness

**DOI:** 10.1101/2024.08.28.610123

**Authors:** Maria Papadaki, Eleftherios Pavlos, Marc Dubourdeau, Vincent Bailif, Kamal Badirou, Ioanna-Evdokia Galani, Dimitris Papelis, Natalia Kamperi, Vassiliki Triantafyllia, Lena Siouti, Maria Salagianni, Maria Manioudaki, Nikolaos Paschalidis, Giannis Vatsellas, Evangelia Koukaki, Vasiliki Rapti, Dimitris Thanos, Nikoletta Rovina, Garyfallia Poulakou, Aurélie Cobat, Jean-Laurent Casanova, Constantin Tamvakopoulos, Evangelos Andreakos

**Author notes:** Laboratory of Immunobiology, Center for Clinical, Experimental Surgery and Translational Research, BRFAA, Athens, Greece. Electronic address.

## Abstract

Severe COVID-19, caused by SARS-CoV-2 infection, is characterized by excessive inflammation leading to the development of pneumonia and acute respiratory distress syndrome. Bioactive lipid mediators (LMs) derived from ω6 and ω3 polyunsaturated fatty acids are central to the regulation of inflammation, controlling both its initiation and resolution. Still, their role in COVID-19 remains underexplored. By employing a holistic approach involving the analysis of white blood cell transcriptomes, targeted lipidomics, cytokine and immune cell profiling, across the spectrum of disease severity groups, including mild non-hospitalized patients and healthy individuals, we now show that LM networks are profoundly altered in COVID-19, correlate with inflammatory patterns, and stratify patients according to disease severity. Central to this are CYP450-derived LMs such as 20-HETE, lipid peroxidation metabolites such as iPF2a-VI, and lipoxygenase-derived LMs such as 12-HETE, all of which are major vasoactive mediators of inflammation. Among them, 20-HETE appears to be a promising prognostic biomarker for ICU admission and a potential therapeutic target for severe COVID-19 disease. Our study thus underscores the significance of LM networks in COVID-19 pathophysiology and sheds light into the broader mechanisms driving viral pneumonia in humans.

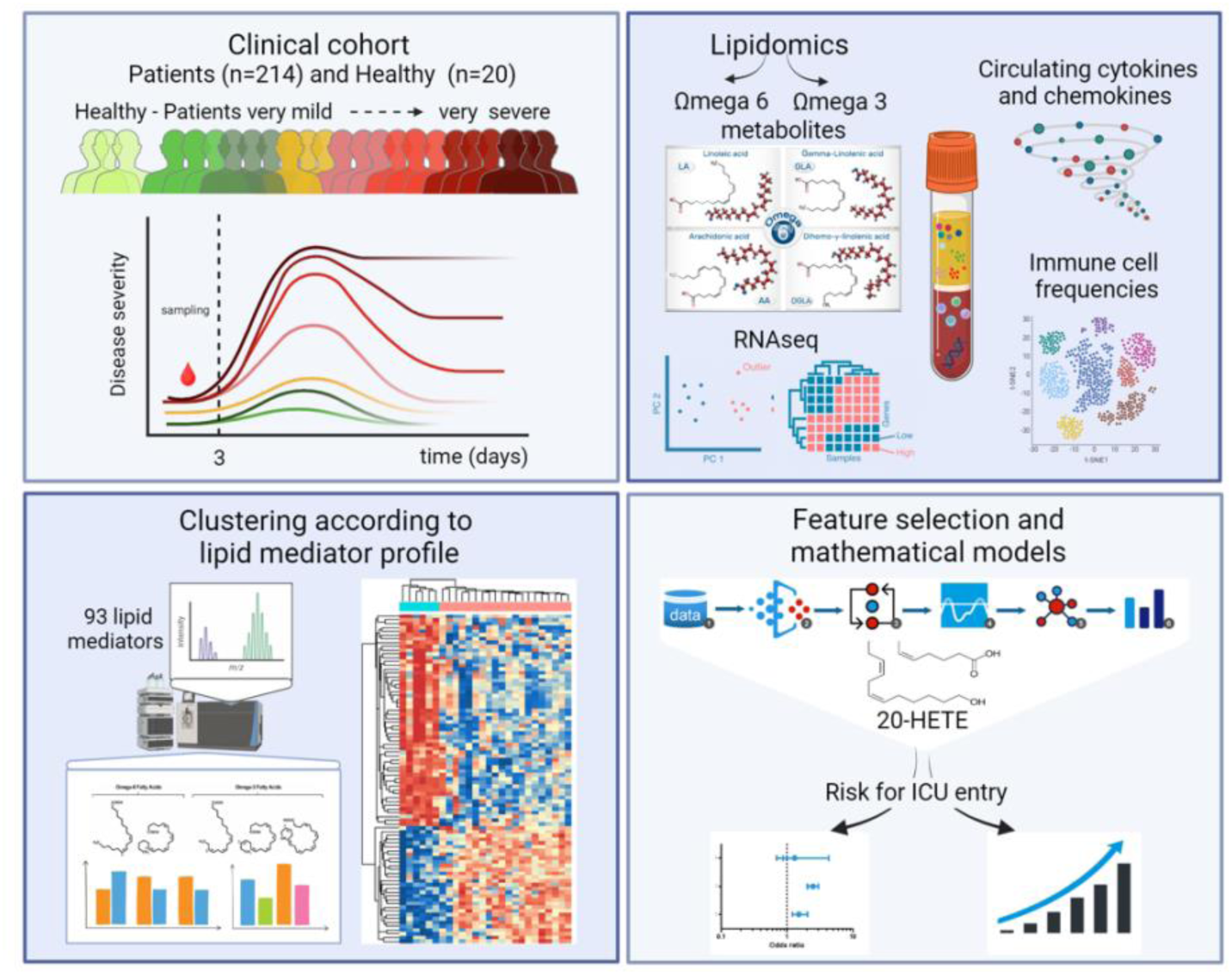

## Introduction

The emergence of coronavirus disease (COVID-19) caused by the novel severe acute respiratory syndrome coronavirus 2 (SARS-CoV-2), has led to a global health crisis with unprecedented rates of morbidity and mortality, and measures of social distancing. Although immunity in the population has increased due to effective vaccines and broad virus exposure, new variants that can avoid pre-existing immunity such as the XBB.1.5, XBB.1.16, JN.1 and the currently and largely dominating BA.2.86 (referred as FLiRT variants) omicron strains are still emerging, emphasizing the need for continuing research on disease pathogenesis and development of better prognostic and therapeutic approaches.

It is well established that dysregulated immunity and hyperinflammation, often occurring as a result of type I IFN deficiencies (genetic or autoimmune) that underlie viral replication and spread, constitute crucial determinants for the development of critical illness (Zhang *et al*., 2020, 2022; Lamers and Haagmans, 2022). Specifically, clinical outcomes and characteristics of critically ill patients are associated with changes in innate and adaptive immunity, including elevated levels of circulating neutrophils (Masso-Silva *et al*., 2022), increased presence of peripherally derived macrophages in the lungs (Merad and Martin, 2020), reduced numbers of circulating T cells (Chen and John Wherry, 2020) and robust cytokine responses (often referred as cytokine ‘storms’), resulting in extensive tissue damage, compromising the gas exchange function and leading to acute respiratory distress syndrome (ARDS) and multi-organ failure (Torres Acosta and Singer, 2020; Lamers and Haagmans, 2022). Bacterial superinfection often occurring during mechanical ventilation of these patients and can further worsen this process (Gao *et al*., 2023).

Although there has been extensive research to identify the various mechanisms leading to hyperinflammation, key immunological effector molecules such as polyunsaturated fatty acid (PUFA)-derived bioactive lipids that participate in the COVID-19 prolonged and uncontrolled state of inflammation has scarcely been studied. Eicosanoids and related metabolites (oxylipins) such as prostaglandins, thromboxanes, leukotrienes and lipoxins are omega-6 lipids formed by oxidation of arachidonic acid (AA) and other PUFAs mainly by 3 enzyme families – cyclooxygenases (COX), lipoxygenases (LOX) and cytochrome P450 (CYP450) (Serhan, 2014). These mostly pro-inflammatory Lipid Mediators (LMs) play a pivotal role in the induction of inflammation and cytokine production (Hajeyah *et al*., 2020). They are critically involved in causing vasodilation, leukocyte extravasation, inflammatory cell activation and prothrombotic changes on endothelial cell function, all highly relevant to COVID-19 (Flaumenhaft, Enjyoji and Schmaier, 2022). Conversely, specialized pro-resolving LMs (SPMs), derived from omega-3 PUFAs such as Docosahexaenoic acid (DHA), Eicosapentaenoic acid (EPA) and n-3 Docosapentaenoic acid (n-3 DPA) including resolvins, protectins and maresins, act to suppress inflammation and inhibit leukocyte extravasation into tissues (Basil and Levy, 2016). They also play a major role in enabling inflamed tissues to return to homeostasis through the coordinated resolution of inflammation, the clearance of cell debris and the downregulation of pro-inflammatory stimulants (Basil and Levy, 2016).

Recent studies have implicated PUFAs in COVID-19. High levels of n-3 PUFAs or a higher n-3:n-6 PUFA ratio have been linked to a lower risk of severe disease *i.e* in patients requiring mechanical ventilation (Mazidimoradi *et al*., 2022; Khan *et al*., 2023). Particularly, the ratio of AA to EPA and n-3 PUFAs, along with IL-6, have been proposed as systemic indicators of poor prognosis, lung damage, and high mortality in COVID-19 (Sertoglu *et al*., 2022). Beyond PUFAs, few studies have provided insight into the profiles of PUFA-derived metabolites or LMs, either pro-inflammatory LMs or SPMs in the peripheral blood (Koenis *et al*., 2021; Palmas *et al*., 2021; Regidor *et al*., 2021; Schwarz *et al*., 2021; Turnbull *et al*., 2022; Irún *et al*., 2023) and lung tissue (Archambault *et al*., 2021; Pérez *et al*., 2022) of COVID-19 patients. Accordingly, increased levels of AA-derived pro-inflammatory lipid mediators (prostaglandins, thromboxanes, and leukotrienes) have been reported in BAL fluid (Archambault *et al*., 2021) and tracheal aspirates (Pérez *et al*., 2022) but also in plasma (Pérez *et al*., 2022) in intubated COVID-19 patients. Notably fewer SPMs (17-HDHA, RvD1, PDX) have been shown to be increased in critically-ill patients when compared to healthy controls (Archambault *et al*., 2021). Interestingly, one study has suggested an important role of a group of seven lipid mediators (TxB4, LTD4, RvE4, 20-COOH-LTB4, 20-OH-MaR1, RvD1, and RvD3) in plasma of severe to critical ill COVID-19 patients that can distinguish patients with from these without mechanical ventilation (Palmas *et al*., 2021). Moreover, other studies have proposed that moderate-to-severe disease is associated with higher levels of COX2 products such as PGE2, PGD2, and PGF2α, while more severe disease is associated with increased ALOX5, ALOX12, and ALOX15 products particularly resolvins, protectins, and lipoxin A4. However, it remains unclear whether these profiles are a driver of severity, a treatment effect (Andreakos, Papadaki and Serhan, 2021), or the result of preexisting conditions, or whether they constitute a hitherto unknown cause of disease susceptibility. It also remains unclear whether these findings still stand in larger or different groups of patients, and groups including patients at all levels of disease severity. Indeed, there have been reports of opposing results with respect to the production of ALOX15-derived SPMs in severe cases compared to mild-to-moderate disease ones (Koenis *et al*., 2021; Schwarz *et al*., 2021). Nevertheless, these studies indicate the potential of using LM profiles to discriminate patients of different disease severity and disease pathophysiology, but also highlight the need for performing more comprehensive analyses involving larger sets of LMs and higher numbers of patients covering the broad disease severity range, as well as combining them with information on established immune-inflammatory mechanisms of the disease.

Here, we performed targeted lipidomics for a total of 93 LMs covering all four bioactive LM metabolomes, and combined them with comprehensive transcriptomic, immune cell profiling and cytokine analyses. We also assessed patient samples across the spectrum of disease severity in order to characterize the complex LM networks that operate during the diverse clinical manifestations of COVID-19, unveil their potential involvement in driving the disease pathophysiology, and relate specific patterns and LMs to risk for worse disease outcomes.

## RESULTS

### Transcriptional deregulation of LM biosynthetic pathways marks critically ill COVID-19 patients

In order to identify molecules and pathways associated with COVID-19 pathophysiology, including biosynthetic processes involved in PUFA-derived LM generation, we first analyzed the white blood cells (WBC) transcriptome of 57 COVID-19 patients, covering the wide spectrum of disease severity, by RNA sequencing (RNAseq). In addition, we analyzed 11 healthy individuals matched for age, sex, and comorbidities. A total of 68 comprehensive RNA-seq gene expression datasets were analyzed (Figure 1).

**Figure 1.**
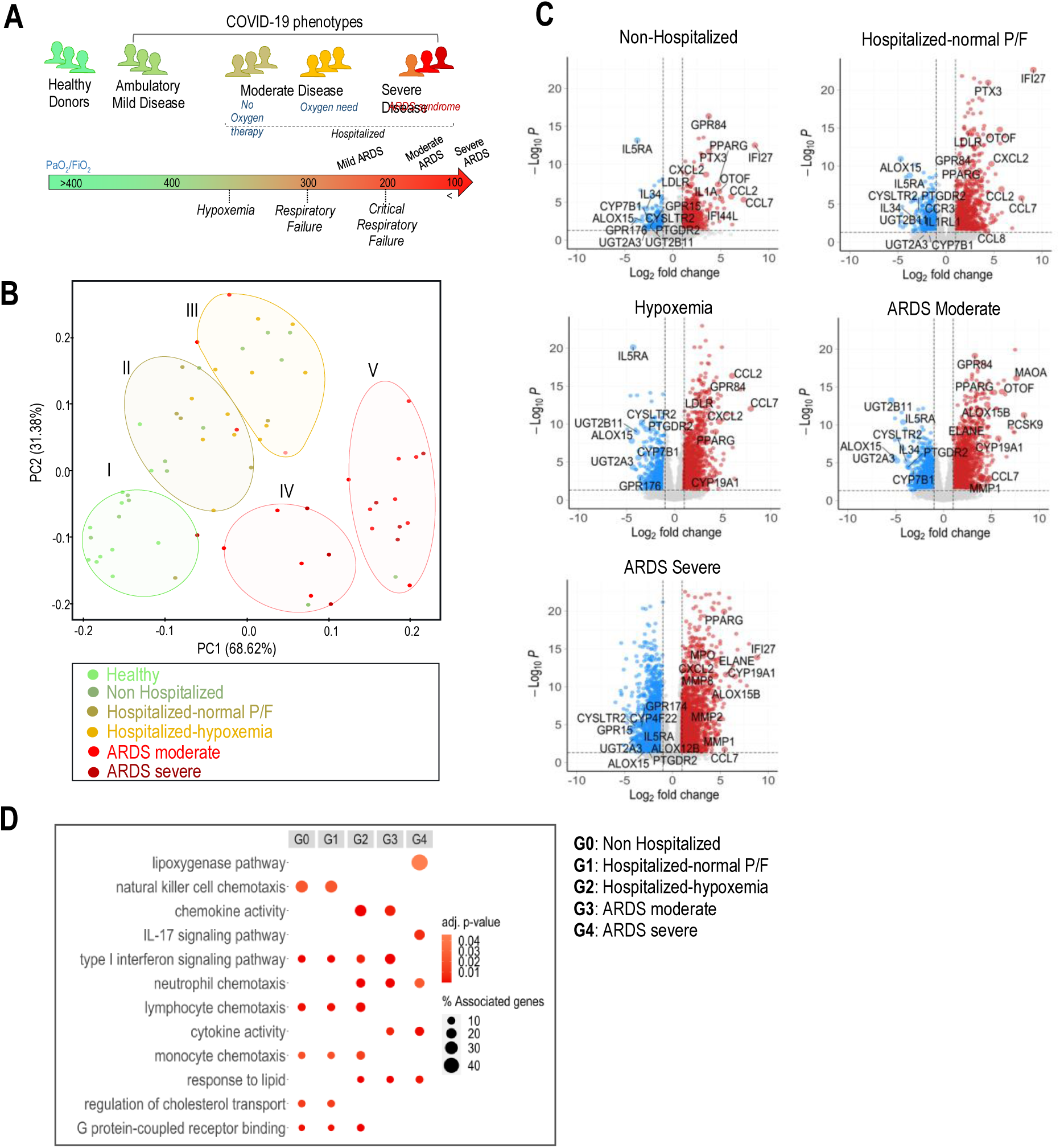
Blood transcriptional signatures of COVID-19 phenotypes reveal dysregulation in lipoxygenase pathway in critical patients. A) Schematic showing the experimental design of COVID-19 patients with different phenotypes. B) Principal-Component Analysis (PCA) of peripheral WBC transcriptomes of non-Hospitalized (*n = 13*), hospitalized with normal PaO2/FiO2 (P/F) (*n=6*), hospitalized with hypoxemia (*n=14*) and ARDS patients: moderate (*n=16*) and severe (*n=8*) and their healthy (n=11) controls. Clusters applied in an unsupervised way after PCA. C) Volcano plots showing the most significantly up/downregulated and relevant DEGs (fc>2, <-2, p<0.05) in each patient category *vs* healthy controls. D) Gene ontology (GO) pathway-enrichment analysis of statistically significant DEGs (p<0.05) of COVID-19 patient subgroups (G0-G4) *vs* healthy controls. Dot size indicates % of associated genes per pathway and color indicates the adjusted p-value per term.

**Table 1.** Demographics and clinical characteristics of the main study patients.

| *Characteristics* |  | HEALTHY | NON-HOSP | HOSP NORMAL | HOSP HYPOX | ARDS MOD | ARDS SEV |
| --- | --- | --- | --- | --- | --- | --- | --- |
| n |  | 19 | 32 | 12 | 25 | 26 | 18 |
| Age (%) | 18-39 years | 2 (10.5) | 9 (28.1) | 5 (41.7) | 3 (12.0) | 2 (7.7) | 1 (5.6) |
|  | 40-59 years | 7 (36.8) | 14 (43.8) | 3 (25.0) | 12 (48.0) | 8 (30.8) | 9 (50.0) |
|  | 60+ years | 10 (52.6) | 3 (9.4) | 4 (33.3) | 10 (40.0) | 16 (61.5) | 8 (44.4) |
|  | unknown | 0 (0.0) | 6 (18.8) | 0 (0.0) | 0 (0.0) | 0 (0.0) | 0 (0.0) |
| Sex (%) | Male | 8 (42.1) | 9 (28.1) | 5 (41.7) | 15 (60.0) | 16 (61.5) | 14 (77.8) |
|  | Female | 11 (57.9) | 23 (71.9) | 7 (58.3) | 10 (40.0) | 10 (38.5) | 4 (22.2) |
| BMI (%) | normal weight | 7 (36.8) | 10 (31.2) | 2 (18.2) | 2 (8.0) | 3 (11.5) | 3 (16.7) |
|  | overweight | 4 (21.1) | 12 (37.5) | 2 (18.2) | 9 (36.0) | 10 (38.5) | 8 (44.4) |
|  | obese | 2 (10.5) | 4 (12.5) | 2 (18.2) | 3 (12.0) | 9 (34.6) | 6 (33.3) |
|  | unknown | 6 (31.6) | 6 (18.8) | 5 (45.5) | 11 (44.0) | 4 (15.4) | 1 (5.6) |
| Ward (%) | YES | 0 (0.0) | 0 (0.0) | 12 (100.0) | 25 (100.0) | 24 (92.3) | 15 (83.3) |
| ICU (%) | YES | 0 (0.0) | 0 (0.0) | 0 (0.0) | 0 (0.0) | 20 (76.9) | 18 (100.0) |
| WHO score (mean (SD)) |  | 0.00 (0.00) | 2.06 (0.25) | 4.33 (0.49) | 4.56 (0.51) | 7.58 (2.00) | 8.44 (1.10) |
| ARDS based on worst P/F (%) | HEALTHY | 19 (100.0) | 0 (0.0) | 0 (0.0) | 0 (0.0) | 0 (0.0) | 0 (0.0) |
|  | NON-HOSP | 0 (0.0) | 32 (100.0) | 0 (0.0) | 0 (0.0) | 0 (0.0) | 0 (0.0) |
|  | HOSP NORMAL | 0 (0.0) | 0 (0.0) | 12 (100.0) | 0 (0.0) | 0 (0.0) | 0 (0.0) |
|  | HOSP HYPOX | 0 (0.0) | 0 (0.0) | 0 (0.0) | 25 (100.0) | 0 (0.0) | 0 (0.0) |
|  | ARDS MOD | 0 (0.0) | 0 (0.0) | 0 (0.0) | 0 (0.0) | 26 (100.0) | 0 (0.0) |
|  | ARDS SEV | 0 (0.0) | 0 (0.0) | 0 (0.0) | 0 (0.0) | 0 (0.0) | 18 (100.0) |
| Timepoint in days (%) | Early (d0-3) | 19 (100.0) | 32 (100.0) | 11 (91.7) | 25 (100.0) | 17 (65.4) | 11 (61.1) |
|  | Late (>d3) | 0 (0.0) | 0 (0.0) | 1 (8.3) | 0 (0.0) | 9 (34.6) | 7 (38.9) |
| Comorbidities: Cardiovascular (%) | YES | 1 (7.7) | 0 (0.0) | 0 (0.0) | 4 (16.0) | 13 (50.0) | 8 (44.4) |
| Comorbidities: Diabetes type II (%) | YES | 0 (0.0) | 1 (3.7) | 1 (8.3) | 4 (16.0) | 10 (38.5) | 4 (22.2) |
| Comorbidities: Dyslipidemia (%) | YES | 3 (23.1) | 1 (3.7) | 1 (8.3) | 5 (20.0) | 6 (23.1) | 5 (27.8) |
| Comorbidities: Arterial hypertension (%) | YES | 0 (0.0) | 2 (7.4) | 3 (25.0) | 7 (28.0) | 7 (26.9) | 8 (44.4) |
| treatment: Corticosteroids (%) | YES | 0 (0.0) | 1 (3.1) | 2 (16.7) | 13 (52.0) | 26 (100.0) | 18 (100.0) |
| treatment: Anakinra (%) | YES | 0 (0.0) | 0 (0.0) | 4 (33.3) | 8 (32.0) | 6 (23.1) | 0 (0.0) |
| treatment: Remdesivir (%) | YES | 0 (0.0) | 0 (0.0) | 8 (66.7) | 16 (64.0) | 15 (57.7) | 10 (55.6) |
| prior treatment: statins (%) | NO | 1 (5.3) | 21 (65.6) | 7 (63.6) | 21 (84.0) | 18 (69.2) | 12 (66.7) |
|  | YES | 0 (0.0) | 0 (0.0) | 3 (27.3) | 4 (16.0) | 8 (30.8) | 6 (33.3) |
|  | unknown | 18 (94.7) | 11 (34.4) | 1 (9.1) | 0 (0.0) | 0 (0.0) | 0 (0.0) |
| Smoking (%) | NO | 18 (94.7) | 18 (56.2) | 4 (33.3) | 16 (64.0) | 16 (61.5) | 11 (61.1) |
|  | YES or ex. | 1 (5.3) | 3 (9.3) | 4 (33.3) | 8 (32.0) | 9 (34.6) | 7 (38.9) |
|  | unknown | 0 (0.0) | 11 (34.4) | 4 (33.3) | 1 (4.0) | 1 (3.8) | 0 (0.0) |
| Onset to Ward [Days (mean (SD))] |  | 0.00 (0.00) | 0.00 (0.00) | 5.58 (2.94) | 7.88 (3.41) | 6.12 (3.58) | 5.00 (1.94) |
| ICU Time [Days (mean (SD))] |  | 0.00 (0.00) | 0.00 (0.00) | 0.00 (0.00) | 0.00 (0.00) | 5.31 (4.69) | 18.39 (18.91) |
| Hospitalization Time [Days (mean (SD))] |  | 0.00 (0.00) | 0.00 (0.00) | 7.75 (2.60) | 9.20 (3.32) | 17.08 (8.94) | 27.83 (17.53) |
| Outcome (%) | Non-hospitalized | 19 (100.0) | 32 (100.0) | 0 (0.0) | 0 (0.0) | 0 (0.0) | 0 (0.0) |
|  | Discharged | 0 (0.0) | 0 (0.0) | 12 (100.0) | 25 (100.0) | 17 (65.4) | 13 (72.2) |
|  | Deceased | 0 (0.0) | 0 (0.0) | 0 (0.0) | 0 (0.0) | 9 (34.6) | 5 (27.8) |

Initially, the overall variance in sample transcriptomes was examined by Principal Component Analysis (PCA). After PCA, a clustering algorithm (k-means) was applied to visualize further patient variability. Results showed a remarkable separation between non-hospitalized (asymptomatic-to-very mild) and moderate-to-critical patients (Figure 1B). Cluster I included mainly healthy (60%) and some very mild (non-hospitalized: 27%) patients, whereas cluster II consisted mainly of very mild to mild patients (non-hospitalized: 25%, hospitalized with normal P/F: 19%, hospitalized with hypoxemia: 38%). Cluster III mostly consisted of hospitalized patients with hypoxemia (53%), and clusters IV and V included 88% and 94% of ARDS patients respectively (Supplementary Table 1). These results demonstrate that samples are clustered according to the severity of the clinical phenotype, suggesting this as the main source of transcriptome variability.

Notably, ARDS patients fell into two separate clusters, cluster IV and cluster V. Although patients in cluster V tended to do better, spending less time in ICU and exhibiting better disease outcomes, while a higher proportion of them was on statin therapy or received dexamethasone (DEX) for a slightly longer time, none of these differences was significant (Supplementary Table 1). Next, volcano plots of Differentially Expressed Genes (DEGs) of distinct patient groups over healthy controls revealed the most differentially regulated genes (Figure 1C). Surprisingly, genes involved in PUFA metabolism, such as those encoding for key enzymes like lipoxygenases (*ALOX15*) and Cytochrome P450 (*CYP7B1*, *CYP4F22*) that biosynthesize various Lipid Mediators (e.g. leukotrienes, prostaglandins and HETEs) and their receptors (*CYSLTR2*, *PTGDR2*, *LGR6*, *GPR176*, *GPR15*), were downregulated in all COVID-19 patients compared to healthy controls (Figure 1C). Furthermore, genes related to glucuronidation reactions, which are involved in PUFA-derived LM degradation and in drug metabolism (Yang *et al*., 2017), such as UGTs (UGT2A3, UGT2B11) were also downregulated in all COVID-19 patients compared to healthy controls (Figure 1C).

Interestingly, *ALOX15*, which is highly expressed in airway epithelial cells catalyzing the conversion of AA to 15-hydroxyeicosatetraenoic acid (15-HETE) (Xu *et al*., 2021), and *UGT2A3 and UGT2B11* which are major contributors to AA metabolites glucuronidation (Turgeon *et al*., 2003), were significantly and consistently downregulated in all patient groups (Figure 1C). This finding strongly suggests a dysregulation of PUFA metabolism in COVID-19 patients, which is intensified with disease severity. On the contrary, among the most regulated genes were genes related to interferon-mediated immune response (*IFI27, OTOF*), lipid metabolism (*PCSK9, OLAH*), and inflammation regulation (*ALOX15B, ALOX12B, MAO-A, CYP19A1, PTX3*). Patients with severe to critical disease (ARDS moderate to severe) showed a significant upregulation of additional genes related to neutrophils activity (*ELANE, MPO, MMPs*), lipoxygenase (*ALOX15B*) and CYP450 (*CYP19A1*) activity (Figure 1C).

Subsequent pathway analysis of DEGs over healthy controls revealed that the lipoxygenase pathway was the most enriched pathway in critically ill patients (Figure 1D), followed by IL-17 signaling, neutrophil chemotaxis, cytokine activity and response to lipid pathways (Figure 1D). When pathway analysis was performed using only the upregulated DEGs, arachidonic acid binding ranked first among the most upregulated pathways (Supplementary Figure 1A), whereas glucuronate metabolic processes were the top downregulated pathway in critically ill patients (Supplementary Figure 1B). These observations are further supported by the selective expression of DEGs related to all lipoxygenases (*ALOX5, ALOX5AP, ALOX12, ALOX12B, ALOX15, ALOX15B, ALOX15P1*), other key LMs enzymes (*PTGS1, PTGS2, TBXAS1, CYP2E1, CYP4F22, CYP4V2, UGT2B11, UGT2A3*) and their receptors (*LTBR4, GPR18, FRP2, CMKLR1*), all of which were also found to be dysregulated among patient groups (Supplementary Figure 2). These findings indicate that dysregulation of lipid mediator metabolism is strongly associated with the severity of COVID-19.

### COVID-19 patients exhibit distinct LM profiles in peripheral blood that relate to disease severity

Having observed at the transcriptional level, a marked dysregulation in the lipoxygenase pathway in severe COVID-19 patients (Figure 1), we evaluated plasma LM profiles in COVID-19 patients with varying disease severity levels. By using a Liquid Chromatography tandem Mass Spectrometry approach (LC-MS/MS), we measured 93 Lipid Mediators (LMs) derived from ω6 and ω3 active metabolomes. To visualize and compare the LM metabolome in every patient, a heatmap was created inputting log-transformed and z-score normalized data on the mean concentration value for each LM measured. Unsupervised hierarchical clustering was therefore applied to test whether LMs could stratify the patients together based on their lipidomic profile. Indeed, LMs were able to categorize patients according to their disease severity. Three major clusters with distinct patterns of LMs were identified (Figure 2). In cluster 1, patients with ambulatory disease were grouped together with healthy (uninfected) individuals (WHO average score of 3, supplementary Table 2A). In Cluster 2, most patients had moderate disease (WHO average score of 5.8), whereas, in Cluster 3, 70% of patients had ARDS (WHO average score of 7.2, Supplementary Table 2A). These results suggest that there is a distinct LM pattern between patient clusters as the disease severity increases.

**Figure 2.**
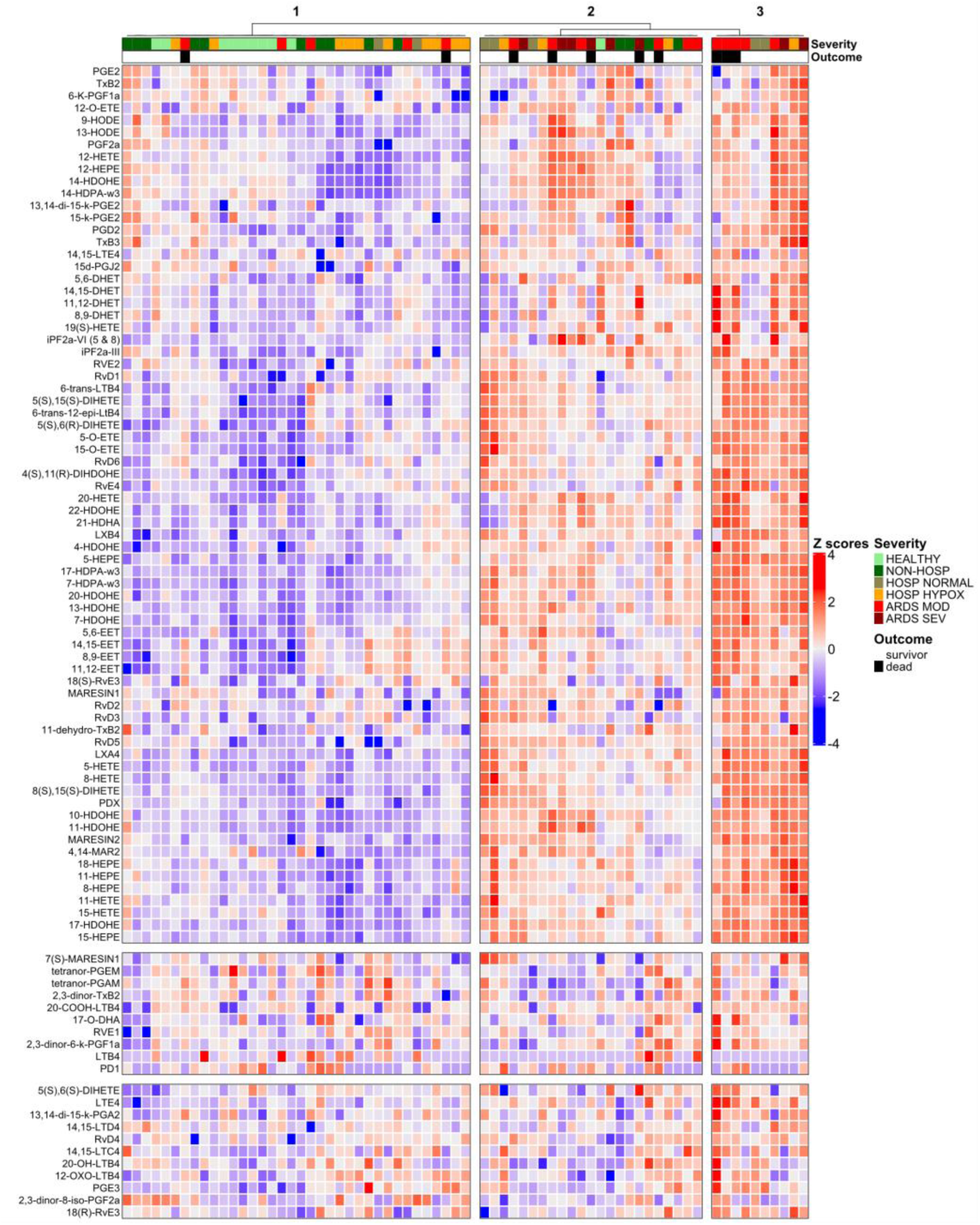
Altered peripheral blood LM profiles across COVID-19 severities. Unsupervised clustered heatmap of 93 LMs in plasma samples collected from COVID-19 patients (n=59) with different disease severities and their healthy controls (HEALTHY, n=10). Each patient column is additionally annotated with data on disease severity (non-hospitalized: NON HOSP, n = 14; hospitalized with normal PaO2/FiO2 (P/F): HOSP NORMAL, n=7); hospitalized with hypoxemia: HOSP HYPOX, n=14; ARDS moderate: ARDS MOD, n=16; ARDS severe: ARDS SEV, n=9), and outcome (survivor or deceased).

Notably, ARDS patients were distributed into two clusters (cluster 2 and 3, Figure 2), with the ones in cluster 3 having a more pronounced LM signature compared to those of cluster 2 (Figure 2, Supplementary Table 2B). However, there were no statistically significant differences between these two clusters of patients in terms of clinical, demographic or other characteristics, although ARDS patients in Cluster 3 had a higher prevalence of diabetes (57% over 17% in cluster 2), and a lower prevalence of hypertension (14% over 50% in cluster 2) (Supplementary Table 2B).

When the distinct LM profiles were analyzed on a per patient group basis according to their absolute amounts, we found that non-hospitalized patients had higher concentrations of COX-derived metabolites of AA such as prostaglandins (i.e 6-k-PGF1a, PGE2, PGD2) and thromboxanes (TXB2) (Supplementary Figure 3), and COX and LOX-derived metabolites of EPA such as resolvins E-series (i.e RvE2) in the blood (Supplementary Figure 4). Mild hospitalized patients, which are only under antibiotic treatment without oxygen support, presented relatively higher levels of LMs and SPMs from all four PUFAs bioactive metabolomes, including higher levels of AA-LOX derived initial metabolites such as 5-HETE and oxo-ETEs, AA-LOX derived SPMs such as lipoxins (LXA4, LXB4), as well as AA-LOX derived leukotrienes (i.e LTB4). In this group, AA CYP450 metabolites, known as nonclassic eicosanoids with mostly anti-inflammatory roles in vascular inflammation (Kim *et al*., 2021) such as EETs (8,9-EETs, 11,12-EETs, 14,15-EETs) were also found increased (Supplementary Figure 3). In terms of n-3 PUFA metabolism, DHA-derived SPMs such as D-series resolvins (RvD1-RvD6) and EPA-derived SPMs such as RvE2 and RvE4 were additionally elevated in this group, indicating a more balanced inflammatory response with strong induction of LMs with potent anti-inflammatory and proresolving roles (Supplementary Figure 4-5). On the other hand, in severe to critical patients increased levels of AA-derived mediators, mostly of pro-inflammatory action, were observed. PGE2, was found increased in our study, which is in line with results from other studies (Koenis *et al*., 2021; Palmas *et al*., 2021). Moreover, ARDS patients exhibited increased levels of a LOX-derived AA metabolite 12-HETE, a CYP450-derived metabolite 20-HETE, and non-enzymatically derived metabolites, such as isoprostanes (iPF2a 5&8). Moreover, autoxidation products of DHA (22-HDOHE, 20-HDOHE, 11-HDOHE, 10-HDOHE, 17-HDOHE, 14-HDOHE) which may serve as markers of oxidative stress (Niki, 2008) have also been found increased in critically ill patients (Supplementary Figure 5).

### Immune-inflammatory parameters are associated with distinct LM profiles of COVID-19 patients and disease phenotypes

We next sought to characterize the immune-inflammatory state of COVID-19 patients, at all levels of disease severity, assess how this relates to PUFA-derived LMs profiles, and determine whether these can be used to discriminate patients. We first performed deep immune phenotyping of cells of the peripheral blood using 30 markers and mass cytometry. We found that there was a gradual increase in neutrophils, classical monocytes and plasmablasts in relation to disease severity, whereas CD4+ and CD8+ T cell populations, regulatory T cells, NK cells, non-classical monocytes, myeloid and plasmacytoid DCs were reduced (Figure 3A-B and Supplementary Figure 6). This was consistent with clinical laboratory tests on cell counts which also revealed increased neutrophil and reduced lymphocyte counts, and an overall increased neutrophil to lymphocyte ratio (Supplementary Figure 7A) as expected (Laing *et al*., 2020). Commonly used clinical biomarkers related to inflammation and tissue damage such as C-reactive protein (CRP), D-dimers and troponin levels were also increased in patients with the most severe disease (Supplementary Figure 7B).

**Figure 3.**
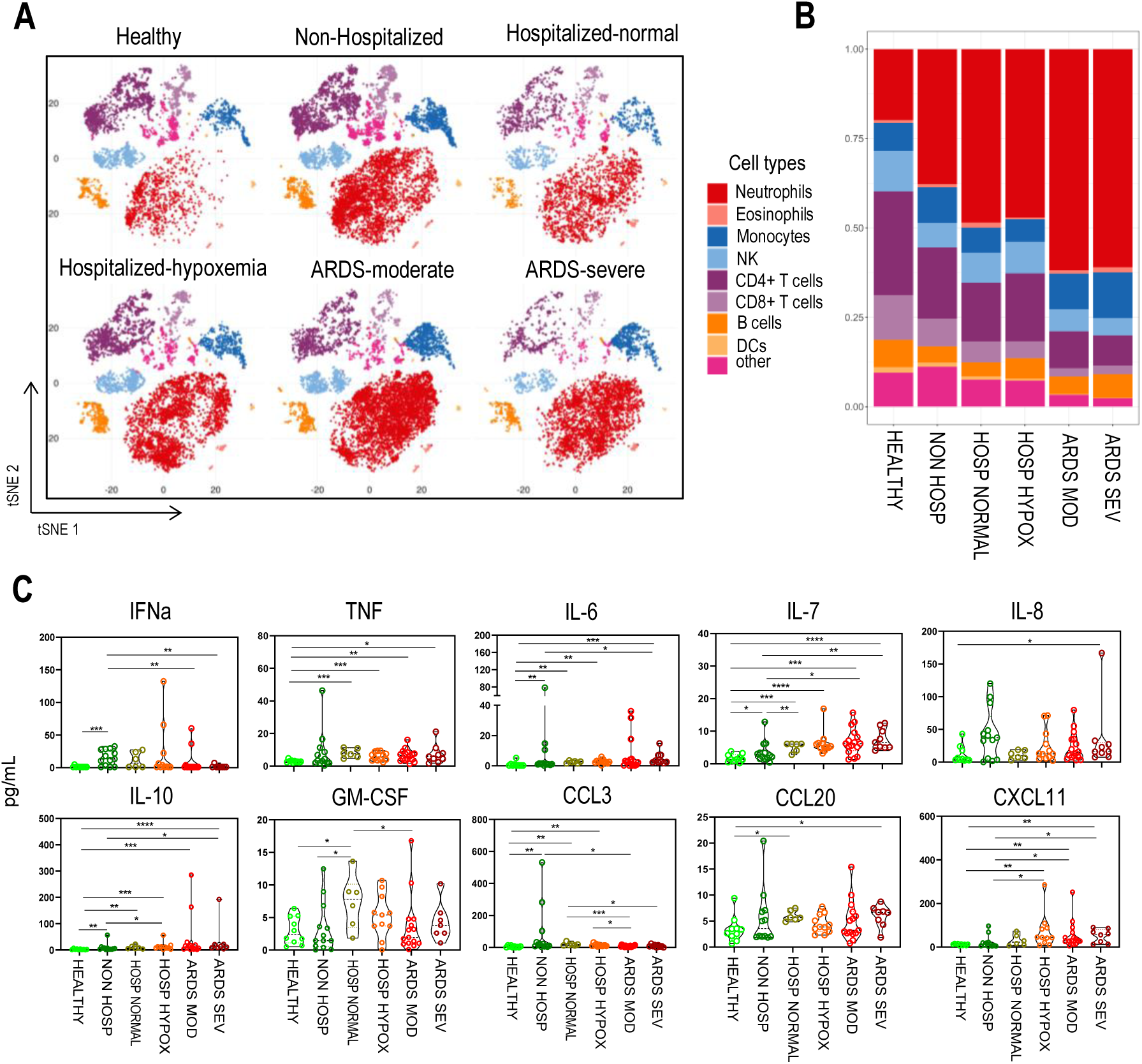
Analysis of inflammatory parameters in COVID-19 patients with different disease severity. A) Cell distribution between patient groups: non-hospitalized (NON HOSP, n=14), hospitalized-normal PaO2/FiO2 (P/F) (HOSP NORMAL, n=7), hospitalized-hypoxemia (HOSP HYPOX, n=14), ARDS moderate (ARDS MOD, n=16), ARDS severe (ARDS SEV, n=9) and their healthy controls (HEALTHY, n=10); visualization with tSNE of CD45^+^ leukocytes. Cells are colored according to cell type classification. B) Stacked barplot represents the mean value of each cell type of tSNE, in each patient group. C) Cytokine levels between patient groups. Data are presented as violin plots with dots showing individual patient measurements, and doted lines median values. *P-*values were determined by Mann Whitney t-test for non-parametric comparisons. \**P* < 0.05, \*\**P* < 0.01 and \*\*\**P* < 0.001.

We then measured the levels of 23 cytokines with high sensitivity multiplex assays in the serum. Molecules such as TNF, IL-6, IL-7, IL-10 and CXCL11/ITAC were significantly higher in COVID-19 patients compared to healthy controls, overall increasing with disease severity but being most significant when mild non-hospitalized patients were compared with the critically ill ones (Figure 3C). In contrast, this pattern was reversed for IFN-α which was significantly elevated only in non-hospitalized patients compared to healthy controls, and had a tendency for higher levels in milder forms of the disease that gradually declined and were almost abrogated in critically ill patients. A similar trend was observed for CCL3/MIP-1a which was higher in non-hospitalized patients but declined thereafter, possibly as a consequence of higher IFN-α levels which strongly induce CCL3/MIP-1a (Salazar-Mather, Lewis and Biron, 2002). For other cytokines and chemokines, there was a variable profile, with IL-8, CCL20/MIP-3a and CX3CL1/Fractalkine being higher in severe critically ill patients, GM-CSF in hospitalized non-hypoxemic patients and CXCL11/ITAC in hospitalized hypoxemic patients (Figure 3C and Supplementary Figure 7C).

Interestingly, when unsupervised clustering based on deep immune cell phenotyping (major cell types) and cytokine data was used (32 parameters in total), patients did not group well according to disease severity (Supplemental Figure 8), possibly reflecting their more variable profile compared to LMs. Thus, although the immune-inflammatory state of patients is linked to the LM profile, the specific immune and inflammatory parameters assessed do not suffice to categorize patients according to disease severity. However, when supervised heat maps for immune cell types and cytokines were added underneath LM profiles, they confirmed that more severe patients that were grouped to clusters 2 and 3 according to their lipidome exhibited an overall higher inflammatory profile compared to milder patients, although several milder patients also had high inflammation (Supplementary Figure 9). This indicates that although higher inflammatory responses are a characteristic of more severe COVID-19 patients, they exhibit a more variable pattern than LMs when patients across the disease spectrum are considered.

### AA-derived CYP450 metabolites such as 20-HETE, iPF2a-VI and 6-trans-12epi-LTB4 discriminate COVID-19 patients at risk of developing critical disease

We next sought to determine whether specific LMs, produced early on in the disease process, can be used to discriminate patients of developing severe disease. For that, we employed feature selection techniques to analyze samples collected within the first three days of hospital admission or visit and identify the most important LMs for that purpose. Among all 93 LMs measured, 9 rated as the most important ones. We found that 6 out of the 9 most important LMs were AA metabolites, either CYP450-derived (20-HETE, 5,6-EET, 5(S),6(R)(R)-DiHET, 6-trans-12epi-LTB4), or non-enzymatically derived (iPF2a-VI, 6-trans-12epi-LTB4), and 3 out of 9 were ω3 DHA LOX-derived metabolites (4S, 11R-DiHDoHE, 7-HDoHE, and RvD5) (Figure 4A). In terms of importance, 20-HETE was ranked first, followed by 6-trans-12epi-LTB4 and iPF2a-VI (Figure 4A). The importance of 20-HETE did not change when inflammatory cytokines and blood cell types were added and feature selection analyses performed in a total of 125 parameters, with some known immune and inflammatory factors such as dendritic cells (DC), T lymphocytes, neutrophils and IL-6 also ranking high (Supplementary Figure 10A).

**Figure 4.**
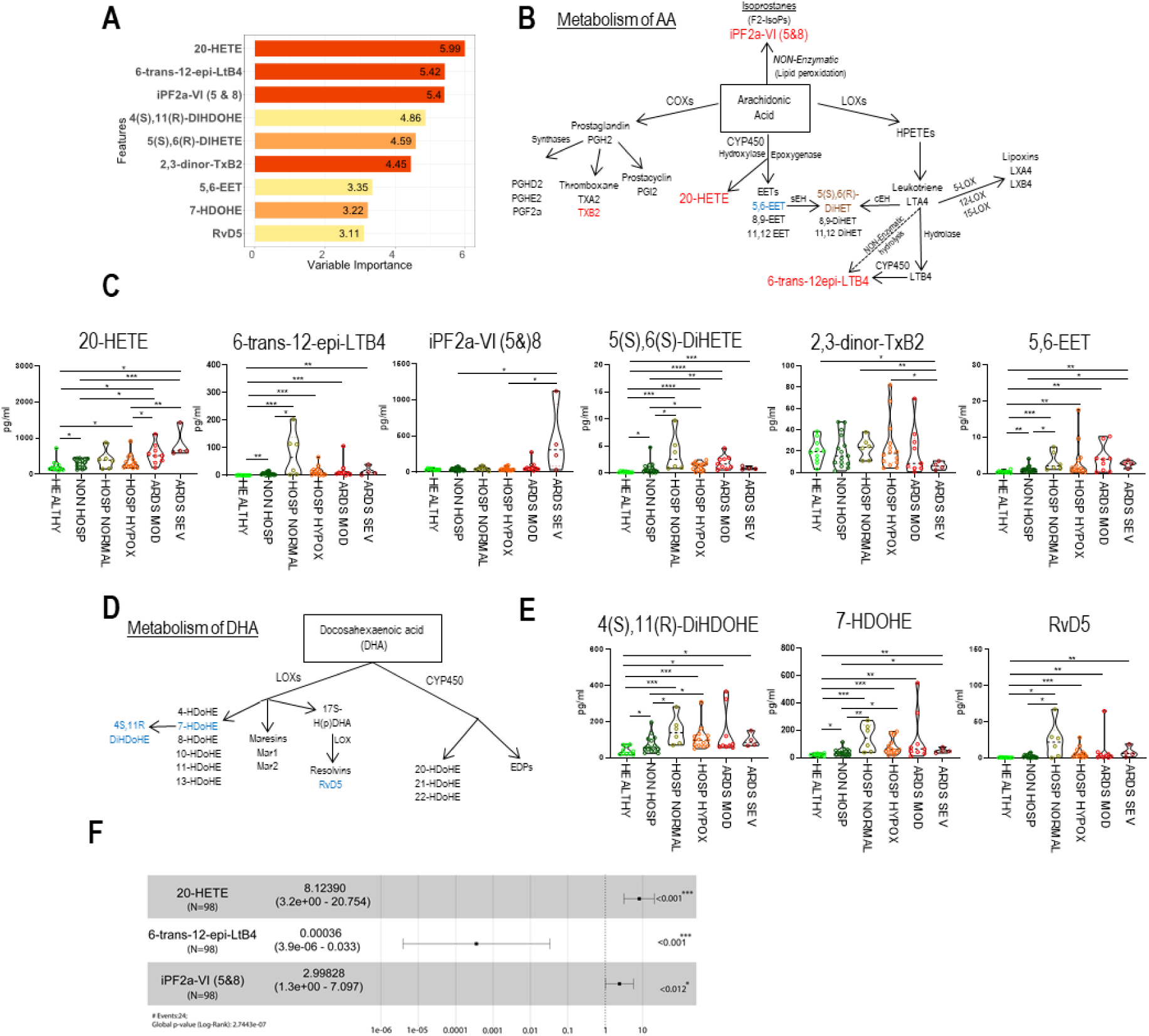
AA-derived CYP450 metabolites have critical role in the pathogenesis of COVID-19. A) Boruta Feature Selection (FS) test performed in patients with an early timepoint only (blood collection at days 0-3 after hospital admission, n=46) and their healthy controls (n=10). B) Schematic representing the most important LMs (in red pro-inflammatory LMs, in blue anti-inflammatory LMs) from Boruta FS test derived from AA metabolism. C) Plasma levels of the most important LMs derived from AA metabolism, of COVID-19 patients with an early timepoint with different severities: non-hospitalized (NON HOSP, n=15), hospitalized-normal PaO2/FiO2 (P/F) (HOSP NORMAL, n=6), hospitalized-hypoxemia (HOSP HYPOX, n=14), ARDS moderate (ARDS MOD, n=9) and ARDS severe (ARDS SEV, n=4) and their healthy controls (HEALTHY, n=10). D) Schematic representing the most important LMs (in red pro-inflammatory LMs, in blue anti-inflammatory LMs) from Boruta FS test derived from DHA metabolism and E) their plasma levels among patient groups. Data are presented as violin plots with dots showing individual patient measurements, and dotted lines median values. *P-*values were determined by Mann Whitney t-test for non-parametric comparisons. \**P* < 0.05, \*\**P* < 0.01 and \*\*\**P* < 0.001. F) Forest plot of top 3 Lipid Mediators with the highest importance score (>5). When Hazard-Ratio (HZ)<1 then the predictor is protective. HZ>1 the predictor is associated with increased risk of ICU admission. More patients with an early timepoint only (blood collection at days 0-3 upon hospital admission, n=98) are added and used for the model.

When LM plasma levels were compared between patient groups, ARDS patients showed increased levels of major AA-CYP450 derived metabolites including 20-HETE (Figure 4B-E). Specifically, in critically ill patients 20-HETE, a well-known pro-inflammatory vasoactive metabolite (Ni and Liu, 2021), was found to be statistically increased, whereas levels of 5,6-EET and its derivative 5(S),6(R)(R)-DiHET that are molecules with opposing anti-inflammatory roles (Ni and Liu, 2021) to 20-HETE were very low (Figure 4C).

Furthermore, it was found that patients with severe ARDS had higher levels of iPF2a-VI molecule which is the most abundant isoprostane product formed during lipid peroxidation and a biomarker of oxidative stress (Niki, 2008) (Figure 4C). Conversely, levels of DHA-LOX derived metabolites with anti-inflammatory effects (Ferreira *et al*., 2022), such as 4(S),11(R)-DiHDOHE, 7-HDOHE and RvD5, were found to be relatively decreased in critically-ill patients when compared to milder cases (Figure 4D-E). Notably, LM levels were more distinct among different patients compared to cytokine levels (Figure 3C), explaining the better discriminatory power LMs exhibit.

Having identified key LMs associated with COVID-19 severity, we next examined whether these can be used to predict disease outcomes. For this reason, a prognostic model for ICU admission was developed using Cox’s regression analysis, based on the most significant LMs identified (importance score >5). Results demonstrated that higher levels of both 20-HETE and iPF2a were found to be strongly associated with increased risk of ICU admission (Figure 4F). By contrast, higher levels of 6-trans-12epi-LTB4 were associated with a reduced risk of ICU admission. This suggests the potential utility of these three LMs as biomarkers in COVID-19 disease outcome.

### Increased 20-HETE levels are associated with higher risk of severe disease independently of vaccination status

As 20-HETE demonstrated notable capacity to predict which COVID-19 patients are at high risk of ICU admission during the early stages of the disease process, we wished to explore that further. One of the main difficulties in identifying robust biomarkers for disease severity in COVID-19 has been the ‘changing’ population with the emergence of new variants and the introduction of vaccination regimens that can alter the immune status of individuals. To account for this, we focused on 20-HETE and developed an additional targeted LC-MS/MS method to measure its levels in more samples and with higher throughput, and applied it to a new patient group that included both non-vaccinated patients as well as fully vaccinated breakthrough patients that have not been investigated before (Figure 5A, Supplementary Table 4). Consistently with the earlier analyses, we found that higher 20-HETE levels in non-vaccinated patients, measured during the first three days of hospital admission, were statistically increased in ARDS patients compared to milder forms of the disease (Figure 5B). Surprisingly, this was also the case in fully vaccinated breakthrough patients, with higher levels of 20-HETE seen in ARDS patients (Figure 5C). Notably, there was no significant increase in 20-HETE levels when non-vaccinated ARDS patients were compared to fully vaccinated ARDS patients (Figure 5D), indicating that vaccination does not change the biomarker potential of 20-HETE for predicting patients at high risk of severe disease and ICU admission.

**Figure 5.**
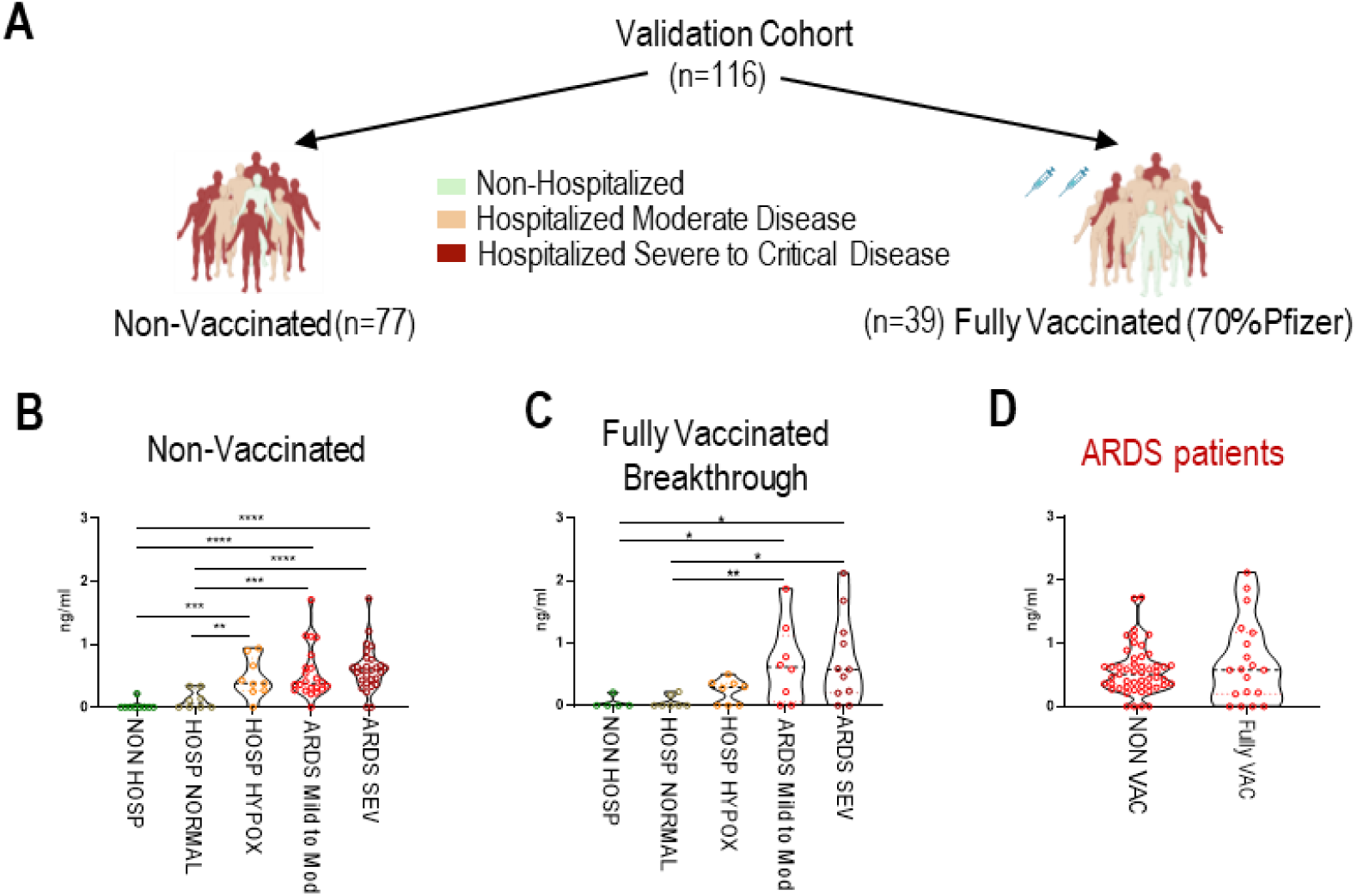
20-HETE plasma levels are increased in critical patients in an independent validation cohort. A) Schematic of validation cohort. 20-HETE clinical validation with LC-MS/MS in a new patient cohort (total, n=116), including B) non-vaccinated patients (n=77) and C) fully vaccinated breakthrough patients (>2 months after 2^nd^ dose, n=39) with different disease severities as in discovery cohort. D) Comparison of non-vaccinated (n=51) and fully vaccinated (n=19) ARDS patients only without any statistical significance. Data are presented as violin plots with dots showing individual patient measurements, and dotted lines median values. *P-*values were determined by Mann Whitney t-test for non-parametric comparisons. \**P* < 0.05, \*\**P* < 0.01 and \*\*\**P* < 0.001.

## Discussion

Most people infected with SARS-CoV-2 will experience mild to moderate respiratory illness and recover without requiring special treatment or hospitalization. However, in some patients SARS-CoV-2 infection will still cause high incidence of severe illness, pneumonia and ARDS, and often despite vaccination. This is due to different reasons. First, susceptibility factors that pre-exist infection, principally age and type I IFN deficiencies of genetic or autoimmune nature, can drive life-threatening COVID-19 (Torres Acosta and Singer, 2020; Manry *et al*., 2022; Zhang *et al*., 2022). Second, mechanisms that come in secondary to disease susceptibility, and that control the immune response and the induction of hyperinflammation, can further worsen the disease. Our study now sheds light into this later process by uncovering novel PUFA-derived LM networks that operate during the development of COVID-19, and that are linked to disease pathophysiology and severity. It also explores their ability to stratify patients according to their clinical phenotype, and identifies novel lipids such as 20-HETE that can be used to predict ICU admission and outcomes. This has profound implications in the way we view the immunopathogenesis of COVID-19, opening up new avenues for the development of novel preventive and therapeutic approaches.

Previous studies have proposed deregulated production of PUFA-derived LMs in COVID-19 patients, with distinct lipid profiles in blood (Koenis *et al*., 2021; Palmas *et al*., 2021; Regidor *et al*., 2021; Schwarz *et al*., 2021; Turnbull *et al*., 2022; Irún *et al*., 2023), immune cells (Koenis *et al*., 2021), and BAL fluid (Archambault *et al*., 2021) linked to increased infection risk and more severe disease. However, specific LMs or LM combinations that can stratify patients and predict outcomes have not been identified as in most cases the patient groups investigated did not include broad disease phenotypes to enable biomarker characterization, while a more limited LM repertoire, mainly focusing on COX and LOX-derived metabolites, was analyzed. Our study thus not only significantly extends this previous knowledge but it also provides a thorough characterization of PUFA-derived LM networks across the spectrum of disease severity, thus enabling the stratification of patients according to their lipidomic profile. It also expands the PUFA-derived LMs studied in the context of COVID-19 by assessing and quantifying 93 molecules covering a broad repertoire of most of the known enzymatic and non-enzymatic derived metabolites of AA, DHA, EPA, linoleic acid and DPA, in patients with different stages of disease severity. Finally, our study assesses LM profiles in relation to immunological information including circulating immune cell populations and cytokines, enabling global and individual interpretation of these profiles at a patient-centered level.

A major observation made early on in our study was that asymptomatic and mild hospitalized patients, rarely explored groups in COVID-19 research, exhibited a more ‘balanced’ response, characterized by increased levels of well-known pro-inflammatory LMs (*i.e* leukotrienes, prostaglandins) as well as anti-inflammatory SPMs (*i.e* Resolvins E-series). On the other hand, severe to critically ill patients had higher levels of more pro-inflammatory and ‘hazardous’ metabolites such as isoprostanes (iPF2a 5&8), autoxidation products of DHA and 20-HETE which was found to be the most important LM related to disease severity among all measured lipids. These findings indicate that specific LM mechanisms are involved in inducing but also limiting inflammation and promoting its resolution, and can be used to discriminate high risk patients according to their lipidome profile with important diagnostic and therapeutic repercussions.

Interestingly, deregulated PUFA-derived LM networks were associated with transcriptional alterations of related pathways in the blood, which were more profound in critically ill patients compared to less severe and milder forms of the disease. There was a dramatic imbalance in the expression of lipoxygenases (*ALOX5, ALOX12, ALOX15*), cyclooxygenases (*PTGS1,PTGS2*) and CYP540 (*CYP2E1, CYP4F22, CYP4V2*) enzymes that are all critically involved in PUFA-derived LM biosynthesis and catabolism (Dyall *et al*., 2022). *ALOX15B*, which is known to be induced in response to hypoxia (Mashima and Okuyama, 2015) was upregulated in ARDS patients, whereas *ALOX15* which suppresses inflammation (Mashima and Okuyama, 2015) was downregulated. Moreover, *CYP2E1*, a monooxygenase that metabolizes AA to 19-HETE that acts as the natural break of the pro-inflammatory 20-HETE (Elshenawy *et al*., 2017), was downregulated. Another CYP450 enzyme *CYP4V2*, which converts FA precursors into n-3 PUFAs, was also found downregulated in severe patients, verifying further the deregulated PUFA metabolism. Impaired expression of several LM receptors (*LTBR4, CMKLR1, GPR18*) was also present in severe patients. These data suggest that deregulated transcriptional pathways related to LM production in the blood are involved in the altered LM patterns observed.

It is well established that severe COVID-19 patients exhibit a hyper-inflammatory response leading to alveolar-capillary damage, pneumonia and ARDS (Zheng *et al*., 2022). In our study, we found that although more severe patients exhibited overall a more inflammatory profile with higher levels of CRP, D-dimers, neutrophils and specific cytokines in the circulation, and lower IFN-α, this was not decisive since significant interindividual variability was observed within the different groups. Classifying, therefore, patients according to the immune cell or cytokine profile, or both, was not as effective, as severe or mild patients did not discriminate well when hierarchical clustering was used for this purpose. It is tempting to speculate that this is related to the crucial role LMs play in vascular homeostasis, controlling vasodilation, leukocyte extravasation and endothelial cell-driven prothrombotic responses, all central to the development of severe COVID-19 (Flaumenhaft, Enjyoji and Schmaier, 2022).

Another major finding of our study was the identification of LMs determining ICU admission risk, with 20-HETE being the most prominent. The prominence of 20-HETE in feature selection analyses remained unchanged even when pro-inflammatory cytokine mediators and immune cell profiling with a well-described role in disease pathophysiology (Costela-Ruiz *et al*., 2020; Gustine and Jones, 2021) were also added, with 20-HETE being also significantly associated with the risk of newly hospitalized patients entering the ICU due to ARDS. Considering the fact that 20-HETE is known to be linked with almost all proposed risk factors for severe COVID-19, such as age, underlying comorbidities *i.e* hypertension, diabetes, obesity, chronic lung diseases, heart, liver and kidney diseases (Elshenawy *et al*., 2017; Gao *et al*., 2021), this makes it an ideal biomarker for disease progression and severity. For this reason, we measured 20-HETE in an independent validation cohort by using a different population including also vaccinated patients. Surprisingly, fully vaccinated critically-ill (breakthrough) patients, presented the same levels of 20-HETE with non-vaccinated ARDS patients. This finding indicates that 20-HETE has the potential of being a prognostic biomarker of critical illness and lung injury broadly applicable to both non-vaccinated and vaccinated COVID-19 patients.

The major link of 20-HETE with disease severity and ICU admission risk can be explained by its multiple and potent roles in the pro-inflammatory and vascular response that drives COVID-19 severity. 20-HETE, produced by CYP450 ω-hydroxylation of AA, is a major vasoactive eicosanoid (Rocic and Schwartzman, 2018) that exhibits diverse effects in inflammation, oxidative stress, endothelial dysfunction and peripheral vascular resistance (Ni and Liu, 2021). Mechanistically, it has been shown to promote vascular inflammation by increasing adhesion molecules (Hoopes *et al*., 2015) and activating nuclear factor-kappa B (NF-κB) (Ishizuka *et al*., 2008) leading to increased production of inflammatory cytokines from activated endothelial cells (Hoopes *et al*., 2015), known to have a crucial role in SARS-CoV-2 infection (Flaumenhaft, Enjyoji and Schmaier, 2022). It has also been demonstrated to drive reactive oxygen species (ROS) production (Medhora *et al*., 2008; Zeng *et al*., 2010; Bou-Fakhredin *et al*., 2021), induce endothelial angiotensin converting enzyme (ACE) expression and activity (Cheng *et al*., 2012), and increase circulating Ang II levels (Sodhi *et al*., 2010) augmenting hypertension and related cardiovascular complications. All these mechanisms are directly related to the vasculopathy that develops in COVID-19 (Flaumenhaft, Enjyoji and Schmaier, 2022). Notably, it is sourced from peripheral blood myeloid and bone marrow-derived cells, including neutrophils, platelets, and endothelial progenitor cells (Tsai *et al*., 2011), all known to be key players in the pathogenesis of COVID-19 (Schulte-Schrepping *et al*., 2020). Interestingly, increased 20-HETE levels have also been observed in other viral and bacterial infections of the respiratory tract (Cuez *et al*., 2010; Tunctan *et al*., 2012; Volzke *et al*., 2020; Schultz *et al*., 2022), suggesting a possibly broader role of this mediator in respiratory infections.

In an effort to understand why 20-HETE is higher in severe COVID-19 patients, and potentially drives disease pathogenesis in that manner, we looked at 20-HETE glucuronidation reactions, that mediate its elimination from the body through kidneys (Yang *et al*., 2017). Remarkably, we found that genes encoding UGTs related to AA-metabolite elimination reactions, were significantly and consistently downregulated among patient groups with the most severe forms of the disease, providing an explanation for the markedly increased levels of 20-HETE observed in these individuals. On the contrary, patients with ambulatory disease did not exhibit significant down-regulation of UGTs, further supporting a milder and more balanced inflammatory response in them.

Our study is not without limitations. First, it needs to be reproduced in larger cohorts with only newly hospitalized patients, of different genetic backgrounds and comorbidities, before 20-HETE can be further explored as a biomarker for disease progression. In addition, it does not establish a cause-effect relationship of 20-HETE and hyperinflammation of COVD-19, although it strongly advocates towards this possibility. Finally, it leaves open the possibility that some cofounding factors such as prior hospitalization treatments can affect these results.

Nevertheless, our study demonstrates the major differences COVID-19 patients with distinct disease severity exhibit in PUFA-derived LM profiles, providing a comprehensive analysis in that respect, as well as their strong association with inflammation and disease outcomes. One of the study’s strengths is its wide range of disease severity groups which provides insights into previously underexplored groups in COVID-19 research, enabling the stratification of patients according to their LM profile. Our study also suggests a critical involvement of eicosanoids in severe COVID-19, and a key role of 20-HETE molecule as a risk factor for critical illness and a potential driver of COVID-19 pathophysiology. Our study, therefore, constitutes a major step forward towards understanding the pathophysiology of COVID-19 and uncovering novel biomarkers and potential mechanisms involved, opening up new avenues for further diagnostic and therapeutic exploration for SARS-CoV-2 but also more broadly other relevant and life-threatening respiratory viral infections.

## Methods

### Study participants

In this noninterventional study, a total of 214 patients including both males and females with different disease severities were used. Study has started with a holistic approach involving the analysis of white blood cell transcriptomes, targeted lipidomics, cytokine and immune cell profiling from 59 COVID-19 participants and 10 Healthy controls. These patients were recruited between 2020 and 2021 from the 1^st^ Respiratory and Critical Care Clinic ward and ICU of the ‘Sotiria’ General Chest Diseases Hospital of Athens, Greece. Healthy, asymptomatic individuals at the time of inclusion served as the control group (n=10-20 Healthy samples). The study conformed to the principles outlined in the Declaration of Helsinki and received approval by the Ethics Committee of the ‘Sotiria’ General Chest Diseases Hospital, Athens, Greece (approval numbers 16707/10-7-18 and 8385/31-3-20). All participants provided written informed consent.

To evaluate the risk of ICU admission, more patients with an early timepoint of hospital admission at sampling (d0-3, n=98) from the same period of pandemic were added for the cox regression Hazard model. We validated our data with another LC-MS/MS method, and a different patient group (n=116), recruited from the beginning of the pandemic until 2022, using also fully vaccinated individuals with breakthrough SARS-CoV2 infection.

The severity of COVID-19 patients was classified based on WHO severity levels (WHO working group, Lancet Infect Dis. 2020). ARDS score has also been added as it is an established lung injury severity marker (JF Murray et al., Am Rev Resir Dis, 1988). Patients are classified into 6 groups based on their worst (Pao_2_/Fio_2_): *Non-Hospitalized*: asymptomatic to very mild non-hospitalized patients with ambulatory disease, with a normal (Pao_2_/Fio_2_); *Hospitalized-normal PaO2/FiO2 (P/F)*: hospitalized patients, with moderate disease but only under antibiotics treatment, with a normal (Pao_2_/Fio_2_); *Hospitalized-hypoxemia*: hospitalized patients with moderate disease, with a Pao_2_/Fio_2_<400; *ARDS mild*: hospitalized patients with severe disease, with a Pao_2_/Fio_2_<300; *ARDS moderate*: hospitalized patients with severe disease, with a Pao_2_/Fio_2_<200; *ARDS severe*: hospitalized patients with severe disease, with a Pao_2_/Fio_2_<100.

### RNAseq analysis

For the analysis of white blood cell transcriptomes, RNA-seq analysis was performed, by purifying total RNA from whole blood leukocytes with the RNeasy Micro kit (QIAGEN). RNA samples were treated with DNase I (QIAGEN) and quantified on a NanoDrop (Thermo Fisher Scientific). Next-generation sequencing libraries were prepared with the TruSeq RNA Library Prep kit v.2 (Illumina) according to the manufacturer’s instructions. Quality of the libraries was validated with an Agilent DNA 1000 kit run on an Agilent 2100 Bioanalyzer.

### Transcriptomics analysis

Samples sequenced on NextSeq 500 (Illumina) were analyzed using standard protocols. Briefly, raw reads were pre-processed using FastQC v.0.11.2 and cutadapt v.1.6, and then mapped to the human genome (GRCh38) using the TopHat version 2.0.13, Bowtie v.1.1.1 and Samtools version v.1.1. The read count table was produced using HTSeq v.0.6. Following filtering of raw read counts with a threshold of 10 in at least one dataset, resulting in a total of 22429 genes. DESEq2 was used to identify the log_2_ fold changes of different groups of patients compared to healthy individuals. DEG transcripts of different COVID-19 patient groups were selected based on an adjusted *P* value cutoff of 0.05 (false discovery rate (FDR) of 5%). Pathway-enrichment analysis was conducted on DEGs over those of healthy individuals using the ClueGO and CluePedia plugin of Cytoscape. TM4 MeV v.4.8 was used for principal component analysis, volcano plots were employed to visualize differentially expressed genes using R package *EnhancedVocano* and pathways plots were generated by utilizing the ggplot2 package in R.

### Lipidomic analysis

A Liquid Chromatography-tandem Mass Spectrometry method was used for the quantification of low-level PUFA metabolites. The extraction protocol and LC-MS/MS analysis were performed by AMBIOTIS SAS (Toulouse, France) using Standard Operating Procedures adapted from Le Faouder et al., 2013. Briefly, Solid phase extraction was used for samples preparation. The method was optimized to obtain a rapid and accurate separation of 93 molecules, with a very high sensitivity of detection and analysis (pg/mL). The lipids were finally eluted with methylformate (MeFor) and methanol (MeOH). After evaporation of the solvent under N2, the residues were recovered in MeOH/H2O and subjected to LC/MS analysis. Analysis was conducted using a scheduled Multiple Reaction Monitoring mode on a 6500 + QTRAP (Sciex) mass spectrometer equipped with an electrospray ionization source in negative mode. The sample was injected beforehand into the Exion LCAD U-HPLC system (Sciex) and eluted on a KINETEX C18 column.

For data visualization, heat maps were performed using *TM4 MeV v.4.8* and *Euclidean* distance was used for hierarchical clustering. Clustering and dendrograms were performed with the *hclust function and ggdendro package*, respectively, in R.

### Validation LC-MS/MS method of 20-HETE

For the quantification of 20-HETE in patient plasma, a liquid chromatography method coupled with mass spectrometry (LC-MS/MS) was developed. Initially, samples were thawed at room temperature and spiked with (internal standard, IS). Protein precipitation was followed by adding cold acetonitrile (ACN). The supernatant was collected and dried at 40°C using a Centrivap vacuum concentrator of Labconco (Kansas city, MO). The residue was reconstituted in 0.1 % Formic Acid and then loaded to the Solid phase extraction (SPE) cartridges. The eluate was evaporated to dryness at 40°C. The remaining residues were reconstituted in 150 μL of mobile phase and transferred into a 96-well plate for LC-MS/MS analysis. The LC-MS/MS method for the quantification of 20-HETE in human plasma was investigated for specificity, linearity, accuracy and precision. The lower limit of quantification (LLOQ) for all analytical runs was set at 0.25 ng/mL. It was defined as the lowest standard on the calibration curve where the analyte peak was at least 5 times the response compared to blank response (S/N > 5) with a precision of RSD≤20 % and an accuracy of 80 – 120%. Briefly, chromatographic separation of 20-HETE and 20-HETE-d6 was carried out using a SunFire C8 column (2.1 x 50 mm, 3.5 μm, Waters, USA) maintained at 40°C and a flow rate of 0.3 mL/min. The optimal LC conditions were as follows: mobile phase A: 90% water, 10% ACN, 0.1% FA, 2 mm ammonium acetate and mobile phase B: 10% water, 90% ACN, 0.1% FA, 2 mm ammonium acetate.

The gradient elution program was as follows: 0-2 min 5%B; 2-8 min 65% B, 8-9 min 80% Β; 9-11 min, 80% Β; 11-12 min 5% Β; 12-15 min 5% B. A SCIEX QTRAP 5500+ (SCIEX, Concord, ON, Canada) was operated in negative ionization mode with multiple reaction monitoring (MRM) transitions of m/z 319.0 → 289.4 and m/z 325.3 → 281.4 for 20-HETE and 20-HETE-d6, respectively. Data acquisition and processing was performed using Analyst software.

### Cytokine analysis

Plasma samples frozen and stored at −80 °C, without other thawing, were analyzed for the presence of IFN-γ, TNF, IL-1β, IL-2, IL-4, IL-6, IL-7, IL-8, IL-10, IL-12 (p70), IL-13, IL-17A, IL-23, CCL3, CCL4 and CX3CL1 with the MILLIPLEX MAP Human High-Sensitivity T cell Panel (Merck Millipore). Thawed plasma aliquots were centrifuged at 13,000 r.p.m. for 10 min at 4 °C immediately before testing. Each assay was performed according to the manufacturer’s protocol for plasma samples, utilizing recommended sample dilutions and standard curve concentrations (Merck Millipore). Samples were analyzed on a Luminex 200 System using Luminex xPonent v.3.1 software according to the manufacturer’s instructions (Merck Millipore). For each cytokine on each assay, the lowest detection limits were in pg ml^−1^: 0.50 for IFN-γ, 0.42 for TNF, 0.2 for IL-1β, 0.24 for IL-2, 0.60 for IL-4, 0.16 for IL-6, 0.33 for IL-7, 0.30 for IL-8, 0.50 for IL-10, 0.24 for IL-12 (p70), 0.20 for IL-13, 0.50 for IL-17A, 8.00 for IL-23, 2.00 for CCL3, 0.80 for CCL4 and 10.00 for CX3CL1.

### CyTOF immunophenotyping

Total leukocytes from patients and healthy donors were isolated and stained with Maxpar Direct Immune Profiling Assay kit (Illumina) of 30 markers, which is optimized for deep immune profiling of human peripheral whole blood by using Mass Cytometry technology (CyTOF). The assay enables comprehensive identification and characterization of 37 immune cell populations, including all major T cell subsets (CD4+ and CD8+ naive, central memory, effector memory, and terminal effector), CD4+ regulatory T cells, CD4–mucosal-associated invariant T cells (MAIT) / natural killer T (NKT) cells, B cell subsets (naive and memory, plasmablasts), natural killer (early and late) cells, T helper (Th) cell phenotypic subsets (Th1-like, Th2-like, and Th17-like), gamma delta (γδ) T cells, monocytes (classical, transitional, and nonclassical), dendritic cell subsets (plasmacytoid and myeloid), granulocytes, basophils, eosinophils, and neutrophils.

### CyTOF data analysis

Flow cytometry standard (FCS) files were normalized using EQ beads and concatenated. Then files were de-barcoded using the barcode key file (Key_Cell-ID_20-Plex_Pd.csv) in the Fluidigm acquisition software (v. 6.7.1014). Clean-up gates for live single cells and elimination of non-cell signals were manually conducted using the web-plat software, Cytobank (v.9.1). Data were analyzed using a previously described R-based pipeline (Nowicka *et al*., 2019). In brief, data were imported and transformed for analysis using the read.flowSet function from the flowCore (v.2.6.0) package (Hahne *et al*., 2009) and the prepData with option (cofactor = 5) function from the CATALYST (v.3.16) package (Nowicka *et al*., 2017), respectively. Clusters were visualized using t-distributed stochastic neighbor embedding (t-SNE) and subsequently annotated based on protein markers expression. Stacked barplot created using ggplot2 R package to visualize cell type abundancies.

### Statistical analysis

The study population’s baseline and admission data were described using descriptive statistics.

Using R package tableone, categorical variables were displayed as numbers and proportions, and continuous variables as means and SD. The ComplexHeatmap package in R was used to construct each heatmap, and Euclidean distance was used for hierarchical clustering. Boruta package in R, a wrapper algorithm around Random Forest was employed to choose the most relevant features from the dataset. To determine the risk factors for hospitalized patients’ ICU admission, Cox regression analysis was employed. Data preparation was carried out using MS Excel, and all analyses were conducted using R-studio version 4.3.0. Statistical significance was set at a P < 0.05.

## Acknowledgements

The authors thank Professor Athanasios Tzioufas for critically reading the manuscript. E.A. is supported by research grants from the European Commission (IMMUNAID, no. 779295, UNDINE, no. 101057100) and the Hellenic Foundation for Research and Innovation (INTERFLU, no. 1574).

## Author contribution

M.P. and E.P. contributed equally to this work. E.A. designed the study. E.K., V.R., N.R. and G.P conducted the clinical studies and medical evaluation of patients. M.P., E.P., V.T., L.S., M.S., E.K., and V.R. performed sample collection and processing. M.D., V.B. and K.B. performed targeted lipidomics. D.P., N.K. and C.T. performed 20-HETE measurements. M.P. and I.E.G performed RNA-seq experiments. G.V., E.P., M.M. and M.P. performed RNAseq analyses. I.E.G., V.T. and E.A. performed cytokine measurements. M.P. and N.P. performed CyTOF immunophenotyping experiments. E.P. performed all bioinformatic analyses and data visualization. M.P., E.P., and E.A. thoroughly analyzed and interpreted the data. D.T., A.C. and J.L.C. interpreted the data and edited the manuscript. M.P. and E.A. wrote the manuscript and supervised the study. All authors critically read and approved the manuscript.

